# CellCountCV – a web-application for accurate cell counting and automated batch processing of microscopy images using fully-convolutional neural networks

**DOI:** 10.1101/867218

**Authors:** Denis Antonets, Nikolai Russkikh, Antoine Sanchez, Victoria Kovalenko, Elvira Bairamova, Dmitry Shtokalo, Sergey Medvedev, Suren Zakian

## Abstract

The *in vitro* cellular models are promising tools for studying normal and pathological conditions. One of their important applications is the development of genetically engineered biosensor systems to investigate the processes occurring in living cells in real time. Today, there are fluorescence protein based sensory systems for detecting various substances in living cells (for example, hydrogen peroxide, ATP, Ca^2+^ etc.) or for detecting processes such as endoplasmic reticulum stress. Such systems help to study mechanisms underlying the pathogenic processes and diseases and for screening potential therapeutic compounds. It is also necessary to develop new tools for processing and analysis of obtained microimages. Here we present our web-application CellCountCV for automation of microscopy cell images analysis which is based on fully-convolutional deep neural networks. This approach can efficiently deal with non-convex overlapping objects, that are virtually inseparable with conventional image processing methods. The cell counts predicted with CellCountCV were very close to expert estimates (the average error rate was < 4%). CellCountCV was used to analyse large series of microscopy images obtained in experimental studies and it was able to demonstrate the endoplasmic reticulum stress development and to catch the dose-dependent effect of tunicamycin.

## INTRODUCTION

A detailed study of normal and pathological processes within the cells is one of the key goals of modern biomedical research. Current methods of microscopic analysis and visualization of cellular structures, organoids and molecules allow to obtain large sets of images that require further processing and analysis. Thus, the development of effective and precise methods of qualitative and quantitative image processing is an urgent task necessary for automation of microscopy images analysis and investigation of various impacts on the cells, including the screening of potential medicinal compounds. These methods should effectively cope with high variance of cell shapes and sizes, and with their tendency to form dense clusters which makes cell counting and segmentation extremely difficult. Besides, such methods, especially those involving machine learning techniques, must be data-efficient since manual labelling of the datasets is a very tedious task.

The endoplasmic reticulum (ER) is a multifunctional cellular compartment responsible for the protein synthesis, folding and processing. Protein folding in the ER may be impaired under different pathological conditions or stimuli. This condition is known as endoplasmic reticulum stress (1). ER stress activates a complex signalling network, referred as the unfolded protein response (UPR). It is known that ER stress is observed in many neurodegenerative diseases (such as amyotrophic lateral sclerosis, Parkinson’s disease, Alzheimer’s disease etc.) and has crucial functions in immunity and inflammation (1, 2). In addition, UPR is also considered as a target in cancer therapy (3).

XBP1 protein is an important component of unfolded protein response. In ER stress, XBP1 mRNA undergoes specific splicing by IRE1α protein. As a result, the active XBP1 protein is formed, which activates the genes necessary to compensate ER stress (4). Currently, these events can be detected with genetically engineered biosensors based on fluorescent proteins, changing their fluorescence intensity, spectral characteristics and/or localization depending on molecular context. This approach allows to visualize the pathological processes occurring in living cells in real time. The specific sensory systems allowing to visualize XBP1 activation *in vitro* and *in vivo* were developed (5, 6). However, the problem of reliable processing and statistical analysis of large sets of microscopy images to assess the ER stress severity and to assess the efficiency of potential therapeutic compounds remained largely unsolved.

The 293A cells have polymorphic non-convex shapes and variable sizes and they tend to form dense clusters, making it difficult to count and segment the cells. The classical image processing techniques (segmentation based on watershed algorithm, different edge-detection and filtration procedures) were found to be insufficient for this task. Here we describe the development of a deep neural network for quantitative analysis of microscopic images of cells, expressing the ER stress biosensor XBP1-TagRFP. Cell counting was realized with an approach based on the paper by Cohen et al. (7) aimed to a very similar research field. The authors proposed a way to count multiple small objects (as compared to the image size) pertaining to a same category using fully-convolutional neural networks (FCNN), which predict the so-called redundant count maps. This approach can efficiently deal with non-convex overlapping objects, that are virtually inseparable when using conventional image processing methods. Besides, using the redundant count maps approach also helps to decrease necessary manual labelling work since weight sharing property makes fully-convolutional networks significantly less hungry for data, and, moreover, with the selected approach each object requires only one-pixel mark. Cell counts predicted with our model were very close to expert estimates. We have also developed a web-application CellCountCV (Cell Counting with Computer Vision) for counting the cells on phase-contrast microscopy images. CellCountCV was used to analyze large series of microscopy images obtained in experimental studies (3880 images from experiment #1 and 5208 images from experiment #2) and it was able to demonstrate endoplasmic reticulum stress development and to catch the dose-dependent effect of tunicamycin. Thus, this approach can be used for automation of both qualitative and quantitative data analysis in biosensor studies and their applications. The usage examples and the statistical analysis details can be found at: https://github.com/denatns/CellCountCV.

## MATERIAL AND METHODS

### Plasmids construction and production

The fragment of XBP1 gene, that encodes the 26 bp intron sensitive to endoplasmic reticulum stress, was amplified with PCR using the following primers: XBP1-cDNA-F 5’-GCGCGGTGCGTAGTCTGGAGC-3’ and XBP1-cDNA-R 5’-GGTATATATGTGGTCAAAACG-3’ from human brain cDNA. The obtained PCR-product was cloned into the pGEM-T Easy vector (Promega). Several plasmid clones were Sanger sequenced and plasmid clones with 26 bp XBP1 intron were found. One clone with spliced variant of XBP1 gene was also found. The obtained plasmid clones (with unspliced and spliced XBP1 variants) were used as matrix for PCR with primers XBP1-S-F 5’-CAAGCTAGCGCCACCATGGACTACAAAGACGATGACGACAAGGAGAAAACTCATGGCCTTGTAGTTGAG-3’ and XBP1-S-R 5’-AATGGTACCCCCTAAGTCAATACCGCCAGAATCC-3’. XBP1-S-F primer contains the FLAG epitope sequence and Kozak consensus sequence. The obtained PCR-product was cloned into the pcDNA 3.1/Hygro (-) (Invitrogen) vector using the restriction enzymes NheI and Acc65I. TagRFP fragment without start codon was obtained by PCR with TagRFP-woATG-F 5’-GT CGGTACCGTGTCTAAGGGCGAAGAGCTG-3’ n BFPX3-R 5’-GCGCTTAAGTTAATTAAGCTTGTGCCCCA-3’ primers and pTagRFP-N vector (Evrogen) as a DNA source. TagRFP fragment was cloned into the pcDNA 3.1/Hygro (-)-XBP1 vector using the restriction enzymes Acc65I and AflII. In results, two plasmid vectors were obtained pCMV-XBP1-TagRFP, that contains unspliced XBP1 fragment, and pCMV-XBP1-TagRFP_intdel, that contains spliced XBP1 version and constitutively expresses TagRFP for using as a control. Transfection grade plasmid DNA were isolated using the PureLink HiPure Plasmid Miniprep Kit (Thermo Fisher Scientific).

### 293A cells transfection

293A cells were transfected using Polyethyleneimine (PEI, Santa Cruz Biotechnology). Cells were seeded at 60%-70% confluency into a well of 6-well plate. 2 μg of plasmid DNA and 14 μg of PEI were mixed into the 280 μl of DMEM without FBS and antibiotics, incubated 10 min, and added dropwise to the cells. Culture medium was changed to a fresh medium after 6 hours of incubation.

### 293A cells cultivation and ER stress induction

293A cells (Invitrogen) were cultivated in a medium consisting of Dulbecco’s modified Eagle’s medium (DMEM)/F12 (1:1), 10% of fetal bovine serum, 200 mM GlutaMAX (Life Technologies) 1% of a penicillin/streptomycin solution (Life Technologies), and 1× non-essential amino acids (Lonza) at 37 °C and 5% CO_2_. In the first series of experiments to induce endoplasmic reticulum stress, tunicamycin (Abcam) was added to the culture medium at a 10 μg/ml concentration, DMSO was added to the control wells at a 1% concentration. In the second series of experiments tunicamycin was added to the different wells of 6-well plate at a 0, 5, 10 and 15 μg/ml concentration, DMSO was added to the control wells at a 0, 0.5, 1 and 1.5% concentration. Tunicamycin and DMSO were added 24h after transfection. The experiments for all concentrations were repeated twice.

### Microscopy images acquisition

For images production was used the Cell-IQ MLF imaging system (CM Technologies) in the Interinstitutional Shared Center of Cell technologies SB RAS. Phase contrast mode was used to obtain cell images. For the detection of TagRFP was used Cy3-C (Ex/Em, 531nm/593nm) filter. In all cases 10× objective was used.

### Dataset preparation

We have used 20 manually labelled images, split into training and validation set with 8:2 ratio. Each image was split into quadrants to reduce GPU memory usage. Simple augmentation, namely rotations on 90, 180 and 270 degrees, was performed to increase the available amount of training data. Finally, the model was tested on 24 images which were not seen by the model during training and that were not used for hyperparameter tuning. The training and validation image sets were preprocessed by human experts who manually labelled the cells with one-pixel marks.

### Neural network model training and cell counting

The model was greatly inspired by Count-Ception network (1) and aimed to predict the redundant count maps. Each output pixel is supposed to count how many objects are present in its receptive field. Therefore, with this approach, any object is counted multiple times and that is why the result is called the “redundant count map”. The model consists of five inception blocks followed by additional convolutional layers. All activations are rectified linear units (ReLU). After each convolutional or inception layer the batch normalization was applied. Weights were initialized with Glorot uniform approach. The overview of neural network architecture is presented on Fig. 1. The model was trained with Adam optimizer with learning rate set to 0.001. Batch size was set to 1. It was trained for 400 epochs and then the model with the lowest validation loss was evaluated on the test set. Fluorescing cells counting was implemented as a simple red channel binarization according to selected red signal intensity level (0-255, by default the threshold was set to 100); the obtained mask was applied to original grayscale image, then the image was processed by the counting model as usual.

**Figure 1.**
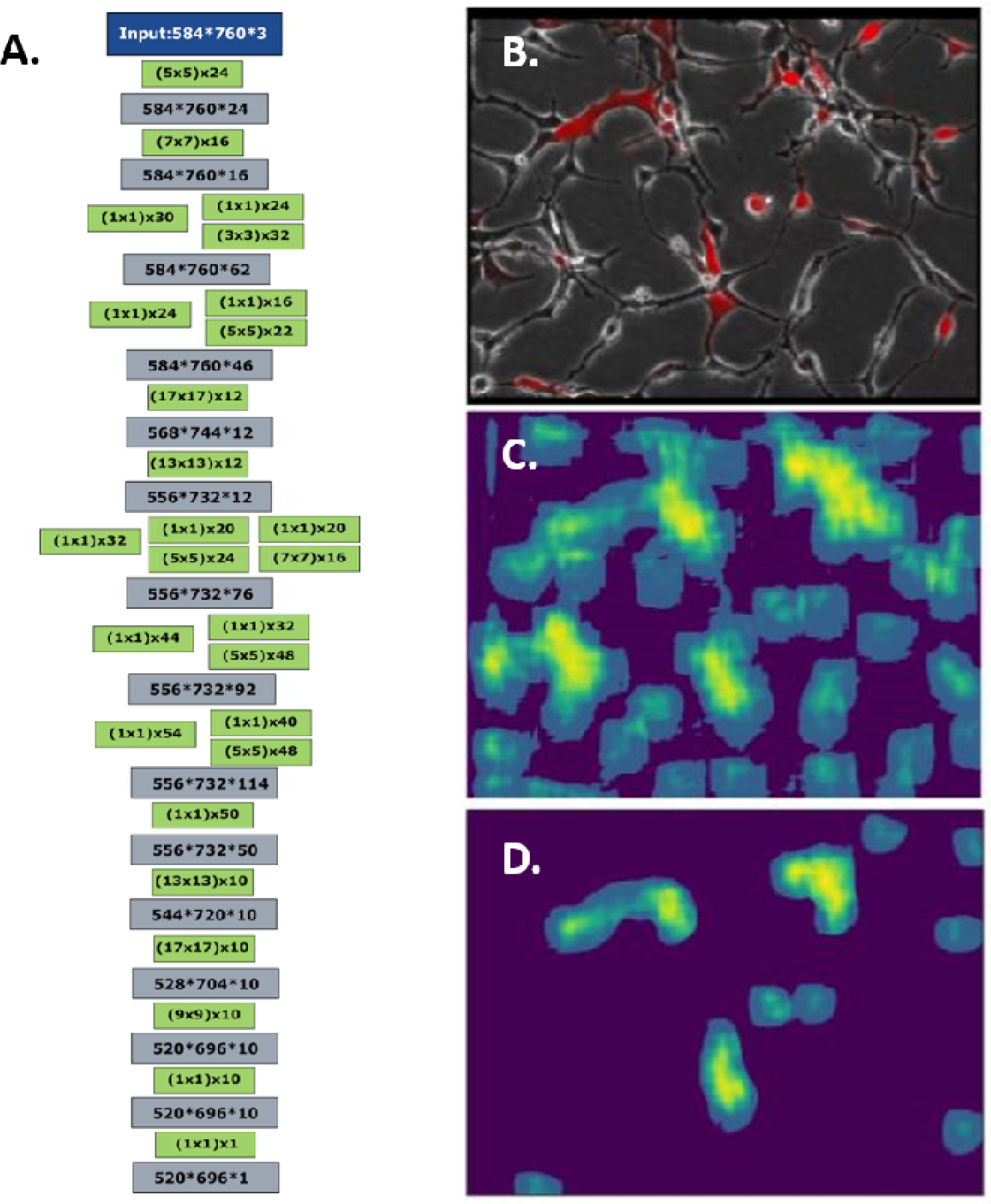
CellCountCV FCNN model architecture and microimages examples. The architecture of developed FCNN model. Green blocks correspond to convolutional layers, where filter sizes are shown in brackets and additional numbers are the numbers of filters. Numbers in grey blocks are output layer sizes (A). Batch-norm blocks are put after each convolutional or inception block and not shown. Examples of original image section (B), corresponding count map (C) and the count map of the cells with red fluorescence above the selected threshold (D).

### Hardware and software

The programs were implemented with Python programming language (v.3.6) using the following libraries: Keras (v.2.1.3) and Tensorflow (v.1.4.1) deep learning packages, that were used to create FCNN model; computer vision and image processing library OpenCV (v.3.1.0) (8) and NumPy package (v.1.14.0) were used for image processing. Web service was created with werkzeug (v.0.14.1) WSGI utility library and json-rpc Python library (v.1.10.3) and run in a docker container created using the official docker image for tensorflow (from https://hub.docker.com/r/tensorflow/tensorflow/) with nvidia-docker plugin to enable GPU usage. API uses JSON-RPC protocol. It has simple graphical user interface implemented with JavaScript library React (v.16.3.1). The model was trained on a workstation running under Ubuntu OS (v.16.04.5) with 32 Gb RAM, Intel^®^ Core™ i7-7740 CPU and 2 Nvidia GTX 1080 GPUs using NVIDIA graphics driver (v.390.87) and CUDA v.9.1.85.

## RESULTS

### Stress induction and other experimental results

In this work, two series of experiments were carried out to induce the ER stress and visualize the activation of the UPR system using a genetically encoded sensor XBP1-TagRFP.

### FCNN model for counting the cells

The network architecture was greatly inspired by the original Count-Ception network described by Cohen et al. (7) which uses 1×1 convolutions. However, the cells and the images used in our study were different from those used to train Count-Ception. The original Count-Ception network was trained and evaluated on VGG, MBM, and Adipocyte cells, which have more rounded shape than 293A HEK cells used here. This increase in complexity made it harder for the original architecture to learn and made us to develop a deeper FCNN architecture. The network structure is shown on Fig. 1A. The two original steps consisting of chained inception layers are also separated by a chokepoint to decrease the size of subsequent hidden layers, and the network also ends with a second chokepoint which restores the size of the output to the of input image. All images used in the current study were 8-bit TIFF files (in either RGB or grayscale), with a size of 1040×1392 px, that is at least 22 times the size of the images been used in the original paper by Cohen et al. (7). As fully convolutional neural network stacks the layers of all size levels of the image it is working with on each level, it requires a vast amount of memory to work with big images. To counter this effect, each individual image was split into quarters and then processed in parallel. The original images have a shape of 1040×1392(×3 for RGB coding), their quarters have a shape of 520×696(×3), and after padding they are resized to 584×760(×3). Each object is originally processed by the model with a 65×65 square kernel of ones. Thus, it is necessary to do image padding to be able to consider the cells located to image borders, so the images were extended by 32 zero pixels from each side. The FCNN thus produces the redundant count map and then the number of the cells is then inferred from the total sum of pixel values of the count-map and scaling it by the number of pixels in a chosen kernel (65×65=4225). This allows also to reduce the total object counting error trough averaging the individual object identification errors by the FCNN. The example of an original image and corresponding count map produced by the model is shown on Fig. 1B.

The training and validation sets were at first labelled together, and then split at 8:2 ratio. Cell culturing and microscopy images acquisition are described in corresponding Materials and Methods sections. To provide more training data, we also used data augmentation: each original image was rotated by 90° 3 times and thus 3 additional images were obtained from each of original ones. The training set series contained 16 images and 4 images were in the validation set. Final performance of our model was assessed with a separate testing set series containing 26 images, none part of which was ever seen by the model or used for hyperparameters tuning.

As there was a very little difference between any pair of successive images, the final performance was evaluated using the images that were the most time-separated from each other. These labelled images were processed with our FCNN model. Then the predicted cell counts were compared to counts provided by human experts.

### Cells counting performance

To assess the performance of produced FCNN model we used the images from two series that were never seen by the model during the training. 26 test images were randomly selected, and then manually labelled by human experts. The total amount of the cells on the test images exceeded 9000. The average error of predicted cell counts for testing images was about 3.12%, which is enough for the method to be considered reliable (Table 1). On 80% of testing images the difference between the manual and predicted cell counts was less than 5%. The highest error percentage of 7.3% was obtained for an image with several clusters of the cells where even the manual counting becomes error-prone. Finally, of all 9247 cells present in the testing dataset, the model counted 9037 of them, thus the overall difference was 2.27%. It indicates that using a set of images (obtained e.g. from different fields of view) will result in higher precision due to averaging the cell counting errors between the neighbour images.

**Table 1.** Validation results obtained on images selected from two series never seen by the model. AE – absolute error, RE – relative error (error percentage).

| Image number | Predicted cell counts | True cell counts | AE | RE (%) |
| --- | --- | --- | --- | --- |
| 1 | 400 | 416 | 16 | 3.85 |
| 2 | 411 | 437 | 26 | 5.95 |
| 3 | 432 | 432 | 0 | 0.00 |
| 4 | 430 | 432 | 2 | 0.46 |
| 5 | 453 | 446 | 7 | 1.57 |
| 6 | 243 | 245 | 2 | 0.82 |
| 7 | 262 | 257 | 5 | 1.95 |
| 8 | 265 | 248 | 17 | 6.85 |
| 9 | 255 | 262 | 7 | 2.67 |
| 10 | 257 | 262 | 5 | 1.91 |
| 11 | 256 | 264 | 8 | 3.03 |
| 12 | 259 | 264 | 5 | 1.89 |
| 13 | 399 | 406 | 7 | 1.72 |
| 14 | 426 | 447 | 21 | 4.70 |
| 15 | 235 | 229 | 6 | 2.62 |
| 16 | 255 | 256 | 1 | 0.39 |
| 17 | 453 | 446 | 7 | 1.57 |
| 18 | 420 | 429 | 9 | 2.10 |
| 19 | 402 | 429 | 27 | 6.29 |
| 20 | 251 | 263 | 12 | 4.56 |
| 21 | 415 | 443 | 28 | 6.32 |
| 22 | 419 | 452 | 33 | 7.30 |
| 23 | 258 | 269 | 11 | 4.09 |
| 24 | 468 | 472 | 4 | 0.85 |
| 25 | 259 | 269 | 10 | 3.72 |
| 26 | 454 | 472 | 18 | 3.81 |
| **Total counts:** | 9037 | 9247 | **Average RE:** | 3.12 |

### Web-application CellCountCV for cell counting

The produced model was used to develop a simple web-application CellCountCV for counting the cells on microscopy images and to count the cells with fluorescence intensity above the user selected threshold. Fluorescing cells counting was implemented using simple binarization mask in image red channel and its application as a filter to a grayscale image, then the image is processed by counting model as usual. Web-service API is implemented with JSON-RPC protocol and we also created simple graphical user interface using React JavaScript library. It accepts a microscopy image and red intensity threshold and returns the cells count and the number of cells with red fluorescence. It was hosted on our institution (IIS SB RAS) server and can be accessed at http://cellcounter.nprog.ru. Besides, it can be used directly through JSON-RPC calls to http://cellcounter.nprog.ru/api. In such a case the image should be encoded as a Base64 string.

### Automation of stress induction analysis in HEK cells with CellCountCV

Images from two ER stress induction time course experiments were processed with CellContCV and the obtained cell counts were analysed. The processing of a single image file on average required about 8.26 seconds on a laptop with NVIDIA GTX 960M GPU and 3.46 seconds on a computing station with NVIDIA GTX 1080 Ti GPUs.

### Endoplasmic reticulum stress induction analysis with CellCountCV

The first series contained 3880 microscopy images (970 images per group with 10 fields of view per time point). This series contained the following groups: Tu – experimental group where ER stress was induced with tunicamycin (10 μg/ml); TagRFP+ – positive control group with constantly expressed TagRFP; DMSO – negative control group of transfected cells with DMSO added (tunicamycin solvent) and Int (intact) – negative control group of transfected cells. The obtained results – cell counts, red cells percentage and red cells percentage change per group per timepoint – are presented on Fig. 2. The original data can be found in supplementary table ST1. Fig. 3 demonstrates the same results but only for the starting and the ending time points.

**Fig. 2.**
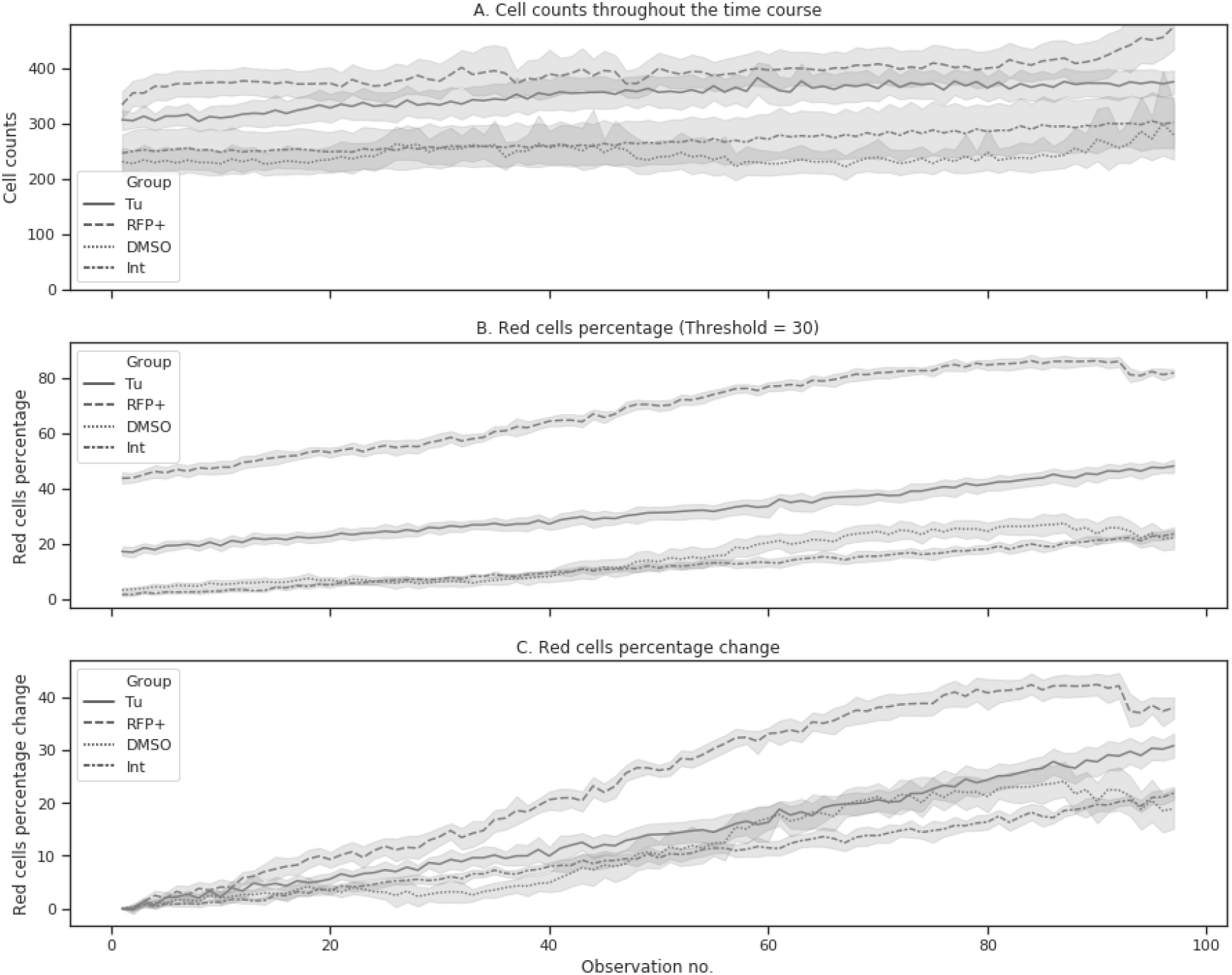
Cell counts and percentage of the red cells during the ER time course. 3880 microscopy images were analyzed (970 images per group with 10 fields of view per time point). Tu – experimental group where ER stress was induced with tunicamycin (10 μg/ml); TagRFP+ – positive control group with constantly expressed TagRFP; DMSO – negative control group of transfected cells with DMSO added (tunicamycin solvent) and Int (intact) – negative control group of transfected cells. (A) CellCountCV was used to estimate the cell counts; (B) red cells were counted at red intensity threshold set to 30 (out of 255) and the percentage of red cells was calculated; (C) the initial red cells percentage was subtracted from all subsequent values for each field of view.

**Fig. 3.**
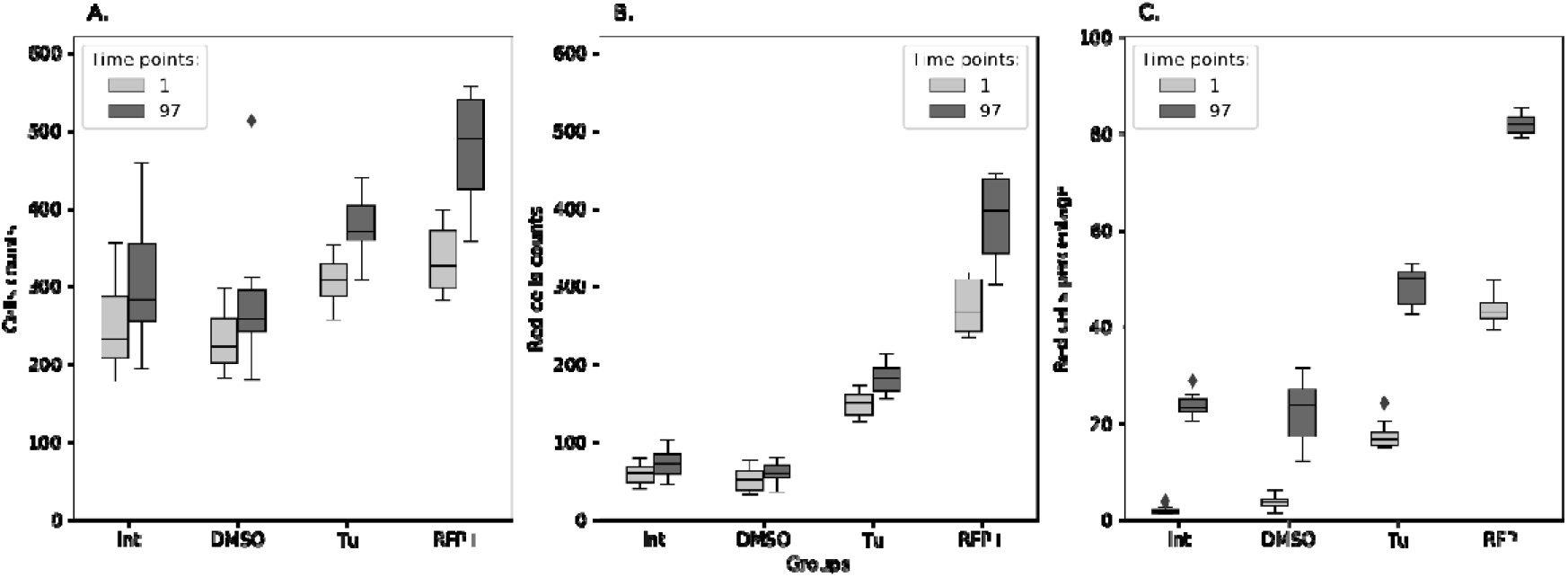
Cell counts (A), red cell counts (B) and red cells percentage (C) in the beginning and at the end of ER time course. Tu – experimental group where ER stress was induced with tunicamycin (10 μg/ml); TagRFP+ – positive control group with constantly expressed TagRFP; DMSO – negative control group of transfected cells with DMSO added (tunicamycin solvent) and Int (intact) – negative control group of transfected cells (N = 10).

The statistical analysis was performed with non-parametric Mann-Whitney test and Bonferroni p-value adjustment for multiple comparisons. Since the initial percentages of red cells were significantly different between the groups (excepting the negative controls) the initial values were subtracted from the subsequent ones. It was found that at the end of experiment the red cell percentages were significantly different between the groups (p < 0.005) excepting negative controls (p = 0.8191) (Table 2).

**Table 2.** Statistical analysis of red cells percentage change at the end of ER time course. The obtained differences in red cells percentages between the starting and the ending time points were compared with non-parametric Mann-Whitney test (two-sided). Bonferroni p-value adjustment was used for multiple testing correction (N = 10).

| Group 1 | Group 2 | U-statistic | p-value | Corrected p-value |
| --- | --- | --- | --- | --- |
| DMSO | Int | 35.0 | 0.1365 | 0.8191 |
| DMSO | TagRFP+ | 0.0 | 0.0001 | 0.0005 |
| DMSO | Tu | 3.0 | 0.0002 | 0.0013 |
| Int | TagRFP+ | 0.0 | 0.0001 | 0.0005 |
| Int | Tu | 0.0 | 0.0001 | 0.0005 |
| TagRFP+ | Tu | 8.0 | 0.0009 | 0.0051 |

Next, we performed the regression analysis using Generalized Estimating Equation (GEE) approach. GEE is a general statistical approach to fit a marginal model for longitudinal/clustered data analysis (9). We used Poisson distribution to model count outcomes – red cells count per 1000 cells – with autoregressive covariance structure. GEE models were built with statsmodels package (v. 0.10.1) for Python. The model demonstrated statistically significant distinctions in all but negative control groups (Int and DMSO) and also demonstrated the significant changes of red cells counts with time (Table 3). The source code used to produce the plots and to perform the analysis can be found https://github.com/denatns/CellCountCV.

**Table 3.** Results of Poisson GEE regression with autoregressive covariance structure. 3880 observations, 40 clusters, cluster size = 97. Scale = 1.0. The model was built with statsmodels package for Python. The basic negative control is the group of intact transfected ER stress-responsive cells (Int), DMSO – is the group of transfected ER stress-responsive cells treated with DMSO; Tu – transfected ER stress-responsive cells treated with tunicamycin and TagRFP+ – positive control – cell constantly producing TagRFP. Time – is a factor of observation time.

| Factor | Coefficient | SD | Z-value | P>\|Z\| | [0.025 | 0.975] |
| --- | --- | --- | --- | --- | --- | --- |
| **Intercept** | 4.4028 | 0.050 | 87.781 | 0.000 | 4.305 | 4.501 |
| **DMSO** | 0.1446 | 0.080 | 1.813 | 0.070 | -0.012 | 0.301 |
| **Tu** | 0.9292 | 0.043 | 21.452 | 0.000 | 0.844 | 1.014 |
| **TagRFP+** | 1.6325 | 0.035 | 46.401 | 0.000 | 1.564 | 1.701 |
| **Time** | 0.0083 | 0.001 | 14.710 | 0.000 | 0.007 | 0.009 |
| **Skew:** | 0.2045 | **Kurtosis:** | -0.2006 |  |  |  |
| **Centered skew:** | -0.4277 | **Centered kurtosis:** | 1.1396 |  |  |  |

### Endoplasmic reticulum stress induction and tunicamycin and DMSO dose-effect analysis with CellCountCV

5208 microscopy images were analysed. A group of cells, constantly expressing TagRFP, was used as a positive control. The cells with TagRFP expression induced under ER stress conditions are referred as experimental group. These two groups were treated with different tunicamycin concentrations. The negative control group were the ER stress-responsive cells treated with different DMSO concentrations (tunicamycin solvent). Cells were counted and the numbers of red cells were obtained with CellCountCV, the percentages of red cells were calculated for each group in 20 fields of view at 24 time points (Fig. 4). Since the initial cell counts and percentages of red cells were significantly different between (and even within) the groups the initial values were subtracted from the subsequent ones, obtained for corresponding fields of view. The changes in total cell counts and red cells percentages throughout the time course are shown on Fig. 5.

**Fig. 4.**
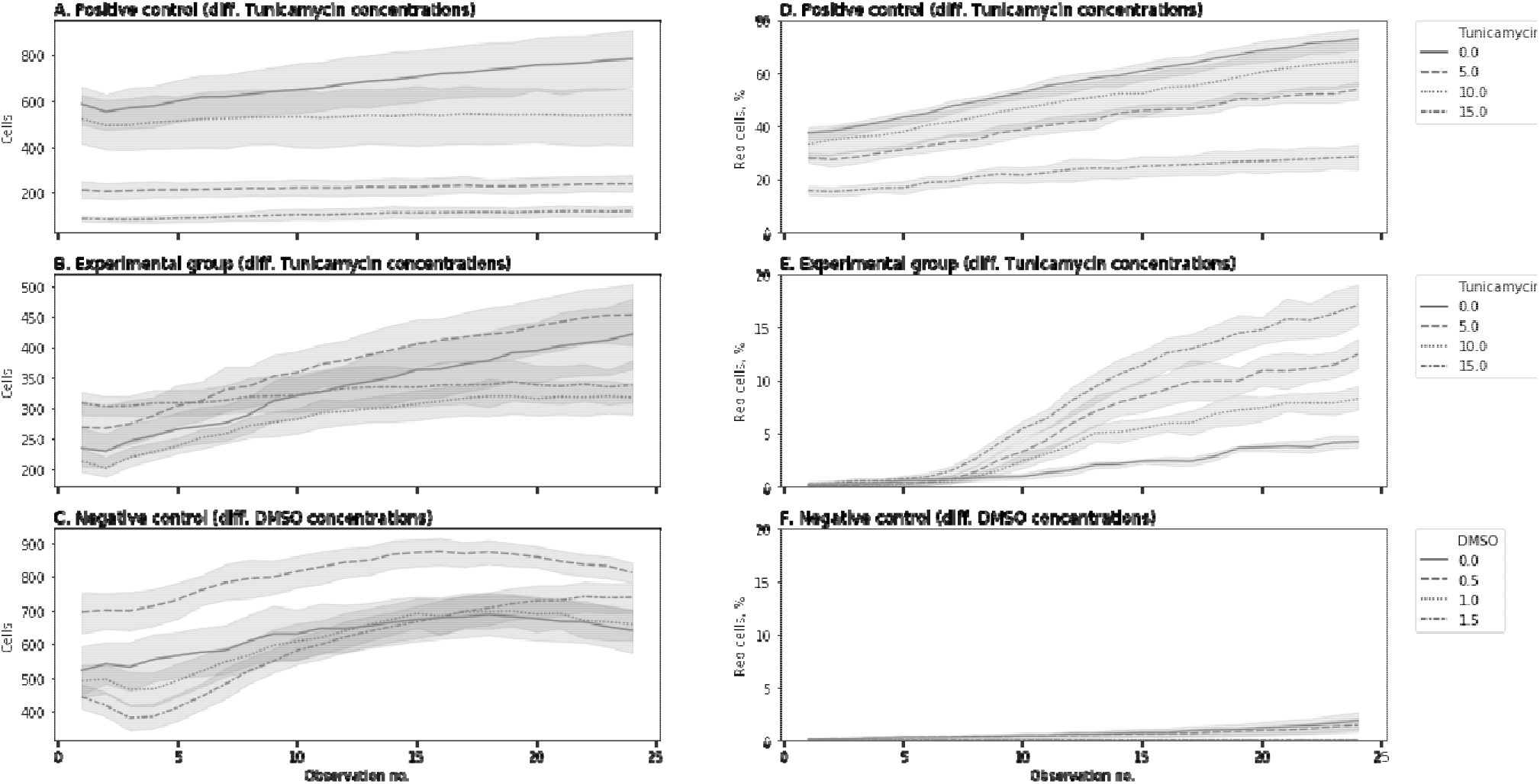
Cell counts and percentage of the red cells during the second ER time course experiment. 5208 microscopy images were analysed. Positive control – a group of cells that constantly expressed TagRFP; experimental group – the cells with TagRFP expression induced under ER stress conditions. These two groups were treated with different tunicamycin concentrations. Negative control group – the ER-responsive cells treated with different DMSO concentrations (tunicamycin solvent). Panels A, B and C show the cell counts throughout the time course and panels D, E and F – the red cells percentage. N = 18.

**Fig. 5.**
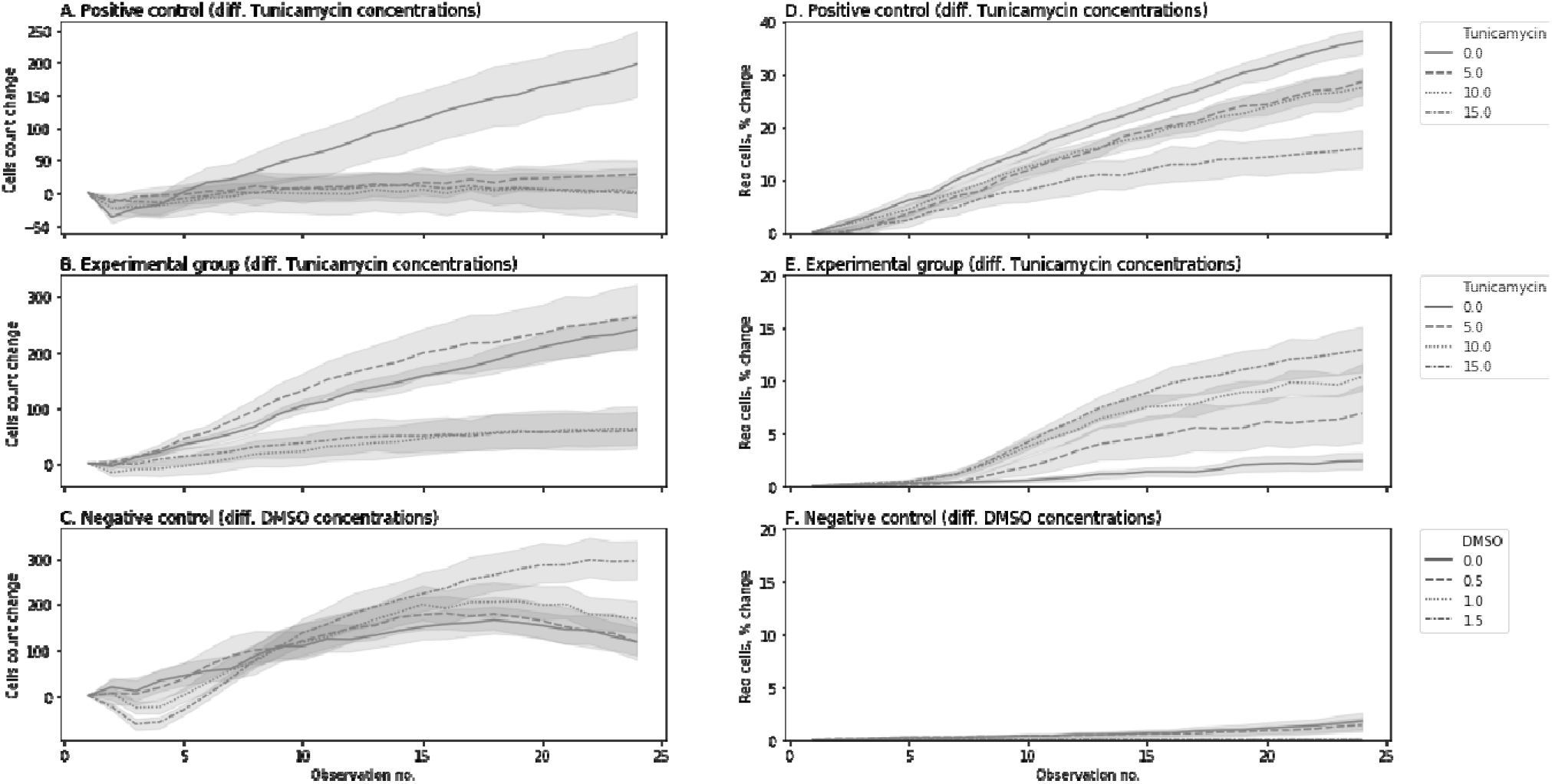
Cell counts and red cells percentage changes during the second ER time course experiment. 5208 microscopy images were analysed. Positive control – a group of cells that constantly expressed TagRFP; experimental group – the cells with TagRFP expression induced under ER stress conditions. These two groups were treated with different tunicamycin concentrations. Negative control group – the ER-responsive cells treated with different DMSO concentrations (tunicamycin solvent). Panels A, B and C show the cell counts change throughout the time course and panels D, E and F – the change of red cells percentage. N = 18.

The statistical analysis was also performed using non-parametric Mann-Whitney test with Bonferroni p-value adjustment. The results of statistical comparison demonstrated significant distinctions between the groups. There were statistically significant distinctions between different tunicamycin and DMSO doses (Table 4). It’s of interest that in positive control group – in cells constantly producing TagRFP – the increased tunicamycin concentration decreased the percentage of red cells percentage while in experimental group tunicamycin dosage had opposite effect (Fig. 6).

**Fig. 6.**
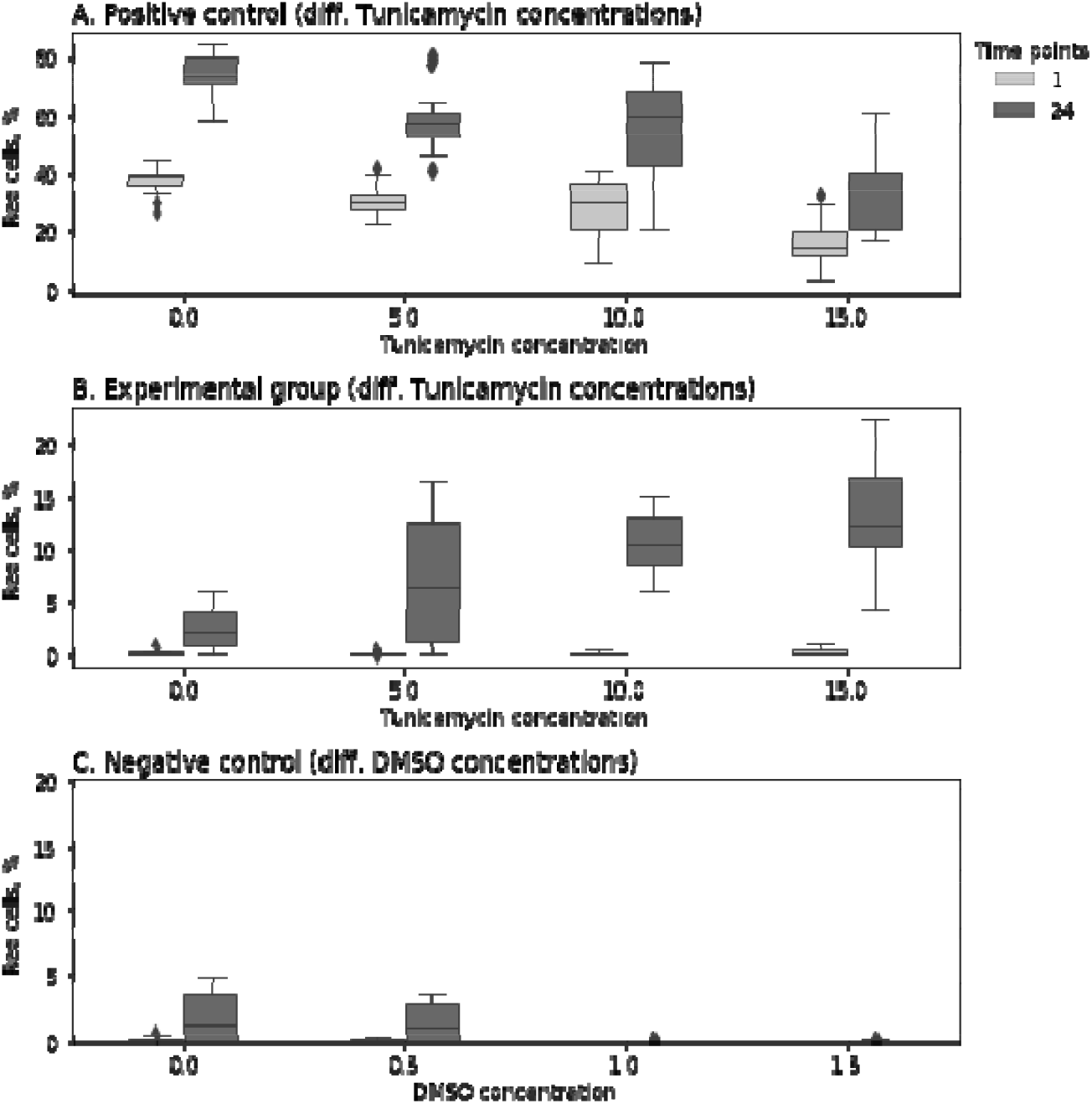
Red cells percentage in the beginning and at the end of the second ER time course. A. – Positive control group with constantly expressed TagRFP treated with different concentrations of tunicamycin; B. – experimental group responsive to ER stress conditions treated with different concentrations of tunicamycin; C. – negative control group treated with different concentrations of DMSO. N = 18.

**Table 4.** Statistical analysis of red cells percentage changes at the end of ER time course between different control and experimental groups, treated with different DMSO and tunicamycin doses. The differences in red cells percentages between the starting and the ending time points were compared with non-parametric Mann-Whitney test (two-sided). Bonferroni p-value adjustment was used for multiple testing correction. N = 18.

| Group 1 | Group 2 | U-statistic | p-value | Corrected p-value |
| --- | --- | --- | --- | --- |
| DMSO_0.0 | DMSO_0.5 | 132.0 | 0.1734 | 1.0 |
| DMSO_0.0 | DMSO_1.0 | 74.0 | 0.0012 | 0.0779 |
| DMSO_0.0 | DMSO_1.5 | 81.0 | 0.0032 | 0.2139 |
| DMSO_0.0 | Exp_Tu_0.0 | 116.0 | 0.0485 | 1.0 |
| DMSO_0.0 | Exp_Tu_10.0 | 0.0 | 0.0 | 0.0 |
| DMSO_0.0 | Exp_Tu_15.0 | 1.0 | 0.0 | 0.0 |
| DMSO_0.0 | Exp_Tu_5.0 | 76.0 | 0.0034 | 0.2212 |
| DMSO_0.0 | Pos_Tu_0.0 | 0.0 | 0.0 | 0.0 |
| DMSO_0.0 | Pos_Tu_10.0 | 0.0 | 0.0 | 0.0 |
| DMSO_0.0 | Pos_Tu_15.0 | 1.0 | 0.0 | 0.0 |
| DMSO_0.0 | Pos_Tu_5.0 | 0.0 | 0.0 | 0.0 |
| DMSO_0.5 | DMSO_1.0 | 83.0 | 0.0043 | 0.2857 |
| DMSO_0.5 | DMSO_1.5 | 90.0 | 0.0094 | 0.6222 |
| DMSO_0.5 | Exp_Tu_0.0 | 99.0 | 0.0149 | 0.9823 |
| DMSO_0.5 | Exp_Tu_10.0 | 0.0 | 0.0 | 0.0 |
| DMSO_0.5 | Exp_Tu_15.0 | 0.0 | 0.0 | 0.0 |
| DMSO_0.5 | Exp_Tu_5.0 | 71.0 | 0.0021 | 0.138 |
| DMSO_0.5 | Pos_Tu_0.0 | 0.0 | 0.0 | 0.0 |
| DMSO_0.5 | Pos_Tu_10.0 | 0.0 | 0.0 | 0.0 |
| DMSO_0.5 | Pos_Tu_15.0 | 0.0 | 0.0 | 0.0 |
| DMSO_0.5 | Pos_Tu_5.0 | 0.0 | 0.0 | 0.0 |
| DMSO_1.0 | DMSO_1.5 | 147.0 | 0.2806 | 1.0 |
| DMSO_1.0 | Exp_Tu_0.0 | 1.0 | 0.0 | 0.0 |
| DMSO_1.0 | Exp_Tu_10.0 | 0.0 | 0.0 | 0.0 |
| DMSO_1.0 | Exp_Tu_15.0 | 0.0 | 0.0 | 0.0 |
| DMSO_1.0 | Exp_Tu_5.0 | 0.0 | 0.0 | 0.0 |
| DMSO_1.0 | Pos_Tu_0.0 | 0.0 | 0.0 | 0.0 |
| DMSO_1.0 | Pos_Tu_10.0 | 0.0 | 0.0 | 0.0 |
| DMSO_1.0 | Pos_Tu_15.0 | 0.0 | 0.0 | 0.0 |
| DMSO_1.0 | Pos_Tu_5.0 | 0.0 | 0.0 | 0.0 |
| DMSO_1.5 | Exp_Tu_0.0 | 1.0 | 0.0 | 0.0 |
| DMSO_1.5 | Exp_Tu_10.0 | 0.0 | 0.0 | 0.0 |
| DMSO_1.5 | Exp_Tu_15.0 | 0.0 | 0.0 | 0.0 |
| DMSO_1.5 | Exp_Tu_5.0 | 1.0 | 0.0 | 0.0 |
| DMSO_1.5 | Pos_Tu_0.0 | 0.0 | 0.0 | 0.0 |
| DMSO_1.5 | Pos_Tu_10.0 | 0.0 | 0.0 | 0.0 |
| DMSO_1.5 | Pos_Tu_15.0 | 0.0 | 0.0 | 0.0 |
| DMSO_1.5 | Pos_Tu_5.0 | 0.0 | 0.0 | 0.0 |
| Exp_Tu_0.0 | Exp_Tu_10.0 | 1.0 | 0.0 | 0.0 |
| Exp_Tu_0.0 | Exp_Tu_15.0 | 3.0 | 0.0 | 0.0 |
| **Exp_Tu_0.0** | Exp_Tu_5.0 | 105.0 | 0.0233 | 1.0 |
| **Exp_Tu_0.0** | Pos_Tu_0.0 | 0.0 | 0.0 | 0.0 |
| **Exp_Tu_0.0** | Pos_Tu_10.0 | 0.0 | 0.0 | 0.0 |
| **Exp_Tu_0.0** | Pos_Tu_15.0 | 4.0 | 0.0 | 0.0 |
| **Exp_Tu_0.0** | Pos_Tu_5.0 | 0.0 | 0.0 | 0.0 |
| **Exp_Tu_10.0** | Exp_Tu_15.0 | 113.0 | 0.0625 | 1.0 |
| **Exp_Tu_10.0** | Exp_Tu_5.0 | 114.0 | 0.0664 | 1.0 |
| **Exp_Tu_10.0** | Pos_Tu_0.0 | 0.0 | 0.0 | 0.0 |
| **Exp_Tu_10.0** | Pos_Tu_10.0 | 7.0 | 0.0 | 0.0 |
| **Exp_Tu_10.0** | Pos_Tu_15.0 | 99.0 | 0.024 | 1.0 |
| **Exp_Tu_10.0** | Pos_Tu_5.0 | 0.0 | 0.0 | 0.0 |
| **Exp_Tu_15.0** | Exp_Tu_5.0 | 79.0 | 0.0045 | 0.2986 |
| **Exp_Tu_15.0** | Pos_Tu_0.0 | 0.0 | 0.0 | 0.0 |
| **Exp_Tu_15.0** | Pos_Tu_10.0 | 21.0 | 0.0 | 0.0003 |
| **Exp_Tu_15.0** | Pos_Tu_15.0 | 126.0 | 0.1307 | 1.0 |
| **Exp_Tu_15.0** | Pos_Tu_5.0 | 3.0 | 0.0 | 0.0 |
| **Exp_Tu_5.0** | Pos_Tu_0.0 | 0.0 | 0.0 | 0.0 |
| **Exp_Tu_5.0** | Pos_Tu_10.0 | 6.0 | 0.0 | 0.0 |
| **Exp_Tu_5.0** | Pos_Tu_15.0 | 66.0 | 0.0013 | 0.083 |
| **Exp_Tu_5.0** | Pos_Tu_5.0 | 0.0 | 0.0 | 0.0 |
| **Pos_Tu_0.0** | Pos_Tu_10.0 | 53.0 | 0.0003 | 0.0197 |
| **Pos_Tu_0.0** | Pos_Tu_15.0 | 2.0 | 0.0 | 0.0 |
| **Pos_Tu_0.0** | Pos_Tu_5.0 | 51.0 | 0.0002 | 0.0156 |
| **Pos_Tu_10.0** | Pos_Tu_15.0 | 45.0 | 0.0001 | 0.0075 |
| **Pos_Tu_10.0** | Pos_Tu_5.0 | 149.0 | 0.3462 | 1.0 |
| **Pos_Tu_15.0** | Pos_Tu_5.0 | 31.0 | 0.0 | 0.0012 |

To investigate the dose-effect of tunicamycin we performed the regression analysis using generalized estimating equation (GEE) approach. Here we also used Poisson distribution to model the count outcome – red cells count per 1000 cells – with autoregressive covariance structure. 3480 observations were used grouped into 145 clusters (according to group, tunicamycin dose and a field of view; cluster sizes were equal to 24 – time points). DMSO treated negative controls were excluded from GEE analysis. The model was built with statsmodels package for Python. The following factors were considered: Tu – tunicamycin concentration, Intron – intron is present in experimental group of cells and is lacking in positive controls, Time – is a factor of observation time. The model demonstrated that all the factors had significant effect and that interactions of Tu and Intron, and Intron and Time were also significant (Table 5). Thus, the regression models also support our conclusions from Mann-Whitney tests about opposite effects of tunicamycin dosage in experimental group and in positive control. The original data can be found in supplementary table ST2. The source code used to produce the plots and to perform the analysis can be found https://github.com/denatns/CellCountCV.

**Table 5.** Results of Poisson GEE regression with autoregressive covariance structure. 3480 observations, 145 clusters, cluster size = 24. Scale = 1.0. DMSO treated negative controls were excluded from GEE analysis. The model was built with statsmodels package for Python. The following factors were considered: Tu – tunicamycin concentration, Intron – intron is present in experimental group of cells and is lacking in positive controls, Time – is a factor of observation time.

| Factor | Coefficient | SD | Z-value | P>\|Z\| | [0.025 | 0.975] |
| --- | --- | --- | --- | --- | --- | --- |
| **Intercept** | 3.9874 | 0.044 | 91.633 | 0.000 | 3.902 | 4.073 |
| **Tu** | -0.0371 | 0.007 | -5.026 | 0.000 | -0.052 | -0.023 |
| **Intron** | -2.5533 | 0.128 | -19.984 | 0.000 | -2.804 | -2.303 |
| **Tu:Intron** | 0.1336 | 0.012 | 11.556 | 0.000 | 0.111 | 0.156 |
| **Time** | 0.0816 | 0.001 | 54.663 | 0.000 | 0.079 | 0.085 |
| **Tu:Time** | -0.0003 | 0.000 | -1.368 | 0.171 | -0.001 | 0.000 |
| **Intron:Time** | 0.0112 | 0.004 | 2.855 | 0.004 | 0.004 | 0.019 |
| **Tu:Intron:Time** | -5.601e-05 | 0.000 | -0.138 | 0.890 | -0.001 | 0.001 |
| **Skew:** | 0.5928 | **Kurtosis:** |  | 0.7584 |  |  |
| **Centered skew:** | -0.6761 | **Centered kurtosis:** |  | 0.9981 |  |  |

## DISCUSSION

With genetic engineering techniques cellular models can be turned into biosensors to visualize cellular stress resulted from particular treatment options. Currently, application of cellular models for studying the mechanisms of different pathogenic conditions and human diseases and developing new efficient therapeutics is actively developing. There is a special interest for cellular models that use the reporter systems based on expression of fluorescent proteins. In such experiments the presence of a fluorescent signal, a change in the spectral characteristics and/or intensity of fluorescence or a change of signal localization within the cell can be detected. To increase the reliability of obtained data, it is necessary to process a large number of microimages. Thus, the researchers have to use either costly proprietary cell-counting systems or semi-automated or manually-operated software packages. This leads to significant time costs and generates the influence of the human factor on the result of the study. We also tried to process these images with either ImageJ (10) or with classical computer vision techniques such as watershed algorithm etc., realized in OpenCV library, but the obtained solutions were found to be inaccurate. That is why we decided to make a dedicated neural network-based model for processing microimages. To develop a prototype of software for automation of endoplasmic reticulum stress assessment we used biosensor XBP1-TagRFP cell line produced from 293A HEK cells.

The developed FCNN model was integrated into a simple web service named CellCountCV. It was found to effectively deal with high variance of cell shapes and sizes and their tendency to form dense clusters. It was also found to be data-efficient as the redundant count maps approach is less hungry for data, and each object requires only one-pixel mark, making the data-labelling less tedious and time consuming. Cell counts predicted with our model were very close to expert estimates. Our model can be used for automation of processing of large sets of microimages. The processing of a single image file on average required about 8.26 seconds on a laptop with NVIDIA GTX 960M GPU, and about 3.46 seconds on a computing station with NVIDIA GTX 1080 Ti GPUs. Thus, the whole set of 5208 images was processed in several hours. If image processing with counting model runs in parallel with image acquisition, then all the results will be ready to the end of the experiment without any significant time delays. The CellContCV application can be used either through simple web user interface at http://cellcounter.nprog.ru or programmatically through JSON-RPC calls to http://cellcounter.nprog.ru/api. Some additional examples and the source code used for plotting and to perform the analysis can be found at https://github.com/denatns/CellCountCV.

The FCNN approach used here also makes it possible to adapt the models for counting any homogenous small objects using the user labelled data. In near future we plan to extend CellCountCV functionality with additional colour filters for fluorescence detection and to create a module for model adaptation to allow researchers to count objects of interest on their images.

Obviously, further development requires the combination of both biosensors and computational model trained on relevant cellular models, for example, on differentiated derivatives of patient-specific induced pluripotent stem cells (11). The induced pluripotent stem cells (iPSC) are widely used in biomedical research and they are especially useful for developing cellular models of human diseases. Such models, based on iPSC, allow to study molecular mechanisms of pathogenesis, to search for target molecules and to screen potential medicinal compounds against human diseases that currently have no efficient therapeutic medications. Such models can also be used to develop personalized and patient-oriented treatment strategies. Here, the 293A HEK cell line was used, that is not relevant to any particular disease. However, this model can be used for primary screening of substances, as it was shown with UPR pathway example.

## AVAILABILITY

CellCountCV web-application can be accessed at http://cellcounter.nprog.ru. Web service can be also accessed programmatically through JSON-RPC calls to http://cellcounter.nprog.ru/api. CellCountCV usage examples, the pretrained FCNN model and the source code used for plotting and for statistical analysis can be found at https://github.com/denatns/CellCountCV.

## ACKNOWLEDGEMENT

The authors are grateful to Dr. Konstantin Orishchenko (Interinstitutional Shared Center of Cell Technologies SB RAS) for help in images production on Cell-IQ MLF imaging system.

## REFERENCES

1. Hetz, C. and Saxena, S. (2017) ER stress and the unfolded protein response in neurodegeneration. Nat. Rev. Neurol., 8, 477–491.

2. Grootjans, J., Kaser, A., Kaufman, R.J. and Blumberg, R.S. (2016) The unfolded protein response in immunity and inflammation. Nat. Rev. Immunol., 8, 469–484.

3. Ojha, R. and Amaravadi, R.K. (2017) Targeting the unfolded protein response in cancer. Pharmacol. Res., 120, 258–266.

4. Yoshida, H., Matsui, T., Yamamoto, A., Okada, T. and Mori, K. (2001) XBP1 mRNA is induced by ATF6 and spliced by IRE1 in response to ER stress to produce a highly active transcription factor. Cell, 107, 881–891.

5. Iwawaki, T., Akai, R., Kohno, K. and Miura, M. (2004) A transgenic mouse model for monitoring endoplasmic reticulum stress. Nat. Med., 1, 98–102.

6. Samali, A., Fitzgerald, U., Deegan, S. and Gupta, S. (2010) Methods for monitoring endoplasmic reticulum stress and the unfolded protein response. Int. J. Cell. Biol., 2010, 830307.

7. Cohen, J.P., Boucher, G., Glastonbury, C.A., Lo, H.Z. and Bengio, Y. (2017) Count-ception: counting by fully convolutional redundant counting. arXiv:1703.08710.

8. Bradski, G. (2000) The OpenCV library. Dr. Dobb’s J. Software Tools, 120, 122–125.

9. Wang, M. (2014) Generalized estimating equations in longitudinal data analysis: a review and recent developments. Adv in Stat., 2014, 1–11.

10. Rueden, C.T., Schindelin, J., Hiner, M.C., DeZonia, B.E., Walter, A.E., Arena, E.T. and Eliceiri, K.W. (2017) ImageJ2: ImageJ for the next generation of scientific image data. BMC Bioinformatics, 18, 529.

11. Ustyantseva, E.I., Medvedev, S.P., Vetchinova, A.S., Minina, J.M., Illarioshkin, S.N. and Zakian S.M. (2019) A platform for studying neurodegeneration mechanisms using genetically encoded biosensors. Biochemistry (Mosc), 84, 299–309.

